# Microbial stimulation of oxytocin release from the intestinal epithelium via secretin signaling

**DOI:** 10.1101/2023.03.09.531917

**Authors:** Heather A. Danhof, Jihwan Lee, Aanchal Thapa, Robert A. Britton, Sara C. Di Rienzi

**Affiliations:** Department of Virology and Microbiology, Baylor College of Medicine, Houston, Texas, USA; Alkek Center for Metagenomics and Microbiome Research, Baylor College of Medicine, Houston, Texas, USA; Department of Neuroscience, Baylor College of Medicine, Houston, Texas, USA; Rice University, Houston, Texas, USA

**Keywords:** *Limosilactobacillus reuteri*, *Lactobacillus reuteri*, oxytocin, secretin, enteroendocrine cells, neuropeptide hormone, gut-brain axis, enteroids, organoids, human, mouse, pig

## Abstract

Intestinal microbes impact the health of the intestine and organs distal to the gut. *Limosilactobacillus reuteri* is a human intestinal microbe that promotes normal gut transit^1^, the anti-inflammatory immune system^2–4^, wound healing^5–7^, normal social behavior in mice^8–10^, and prevents bone reabsorption^11–17^. Each of these functions is impacted by oxytocin^18–22^, and oxytocin signaling is required for *L. reuteri-*mediated wound healing^5^ and social behavior^9^; however, the initiating events in the gut that lead to oxytocin stimulation and related beneficial functions remain unknown. Here we found evolutionarily conserved oxytocin production in the intestinal epithelium through analysis of single-cell RNA-Seq datasets and imaging of human and mouse intestinal tissues. Moreover, human intestinal organoids produce oxytocin, demonstrating that the intestinal epithelium is sufficient to produce oxytocin. We subsequently found that *L. reuteri* facilitates oxytocin secretion directly from human intestinal tissue and human intestinal organoids. Finally, we demonstrate that stimulation of oxytocin secretion by *L. reuteri* is dependent on the gut hormone secretin, which is produced in enteroendocrine cells^23^, while oxytocin itself is produced in enterocytes. Altogether, this work demonstrates that oxytocin is produced and secreted from enterocytes in the intestinal epithelium in response to secretin stimulated by *L. reuteri*. This work thereby identifies oxytocin as an intestinal hormone and provides mechanistic insight into avenues by which gut microbes promote host health.

## Introduction

The microbiome era of biology has heralded a renewed and strengthened understanding that host-microbe interactions in the gut affect not just gut health but also total body health. This knowledge leaves us with the potential capacity to regulate the physiology of multiple organs through manipulation of gut microbial functions. However, achieving this goal is challenged by our limited mechanistic knowledge of how gut microbes interact with their host.

The human intestinal microbe *Limosilactobacillus reuteri* 6475 is one such microbe that can influence the function of multiple host organ systems. *L. reuteri* 6475 reduces gut inflammation in children^24^, adults^25^, and rodent models of chemically induced gut inflammation^2,26–28^, suppresses bone loss in six different murine models of osteoporosis^11–16^ and in a clinical trial of post-menopausal women^17^, promotes skin wound healing in mice^5^ and in humans^6^, and promotes social behavior in six mouse models of autism spectrum disorder^8–10^.

*L. reuteri*’s ability to promote social behavior and wound healing has been demonstrated to require oxytocin signaling^5,9^. Oxytocin is a 9 amino acid peptide hormone produced predominantly in the paraventricular and supraoptic nuclei of the hypothalamus^31^, a brain region involved in the regulation of feeding and social behavior, but also in other organs throughout the body including the enteric nervous system^32^. Oxytocin is a multi-functional hormone that affects not only social behavior^18^, bone^21^, and skin^22^, but also gut motility^19^, inflammation^19,20^, and the intestinal epithelial barrier^19^ (**Table 1**). Therefore, *L. reuteri* signaling through oxytocin could potentially explain many of the systemic effects of this microbe.

**Table 1.** Oxytocin as an effector molecule of *L. reuteri*’s known health benefits.

|  |  |
| --- | --- |
| **Oxytocin function** | **Condition *L. reuteri* affects** |
| Social bonding<sup>18</sup> | Social behavior in autism<sup>8–10*</sup> |
| Anti-inflammatory<sup>19,20</sup> | Inflammation<sup>1–4</sup> |
| Promotes osteoblasts and inhibits bone reabsorption<sup>21</sup> | Osteoporosis<sup>11–17,29</sup> |
| Promotes hair and skin growth<sup>22</sup> | Wound healing<sup>5–7*</sup> |
| Slows gut transit<sup>19</sup> | Gut transit<sup>1</sup> |
| Improves gut barrier integrity<sup>19</sup> | Gut permeability<sup>30</sup> |
| *Demonstrated signaling through oxytocin is required for effect. |  |

Nevertheless, a question remains of how *L. reuteri* impacts oxytocin signaling. *L. reuteri* is believed to predominantly inhabit the human small intestine^33^ and appears to have co-evolved with humans and its other mammalian hosts^1^. Its signaling to hypothalamic oxytocinergic loci requires an intact vagal nerve, implying a signal is sent from the gut to the brain^5,9^. The origins of this signal in the gut, however, remain unknown.

Here, we addressed this question by probing the secreted products of the intestinal epithelium. In doing so, we discovered oxytocin itself is present in the intestinal epithelium in enterocytes. Further, we observed *L. reuteri* can stimulate the secretion of oxytocin and does so through release of secretin, a small intestinal hormone. This work thus advances our understanding of the mechanism by which *L. reuteri* impacts physiology and identifies oxytocin as a previously unrecognized gut epithelial hormone.

## Results

### Oxytocin is produced in the intestinal epithelium

Given the links between *L. reuteri* and oxytocin signaling and the overlap between hormones produced in the gut and brain, we postulated that oxytocin might be present in the gut epithelium. We investigated this possibility by analyzing publicly available mammalian single-cell RNA-Seq (scRNA-Seq) datasets on the intestine. Consistent with our hypothesis, we found an evolutionarily conserved signature of *OXT* transcripts in epithelial cells of mice, macaques, and humans^34–39^ (**Supplementary Table 1**). We subsequently performed a detailed analysis of the human datasets available in the Human Cell Landscape^39^ and Gut Cell Atlas^35^, which reported scRNA-Seq data from multiple regions of the adult intestinal tract. We observed the greatest number of *OXT* positive cells and the greatest expression of *OXT* in the jejunum of the human small intestine (**Fig. 1a, Extended Data Fig. 1a**). Additionally, we found *OXT* expressed in human and mouse organoids derived from the small intestine and colon^37,38,40–42^ (**Supplementary Table 1**).

**Fig. 1:**
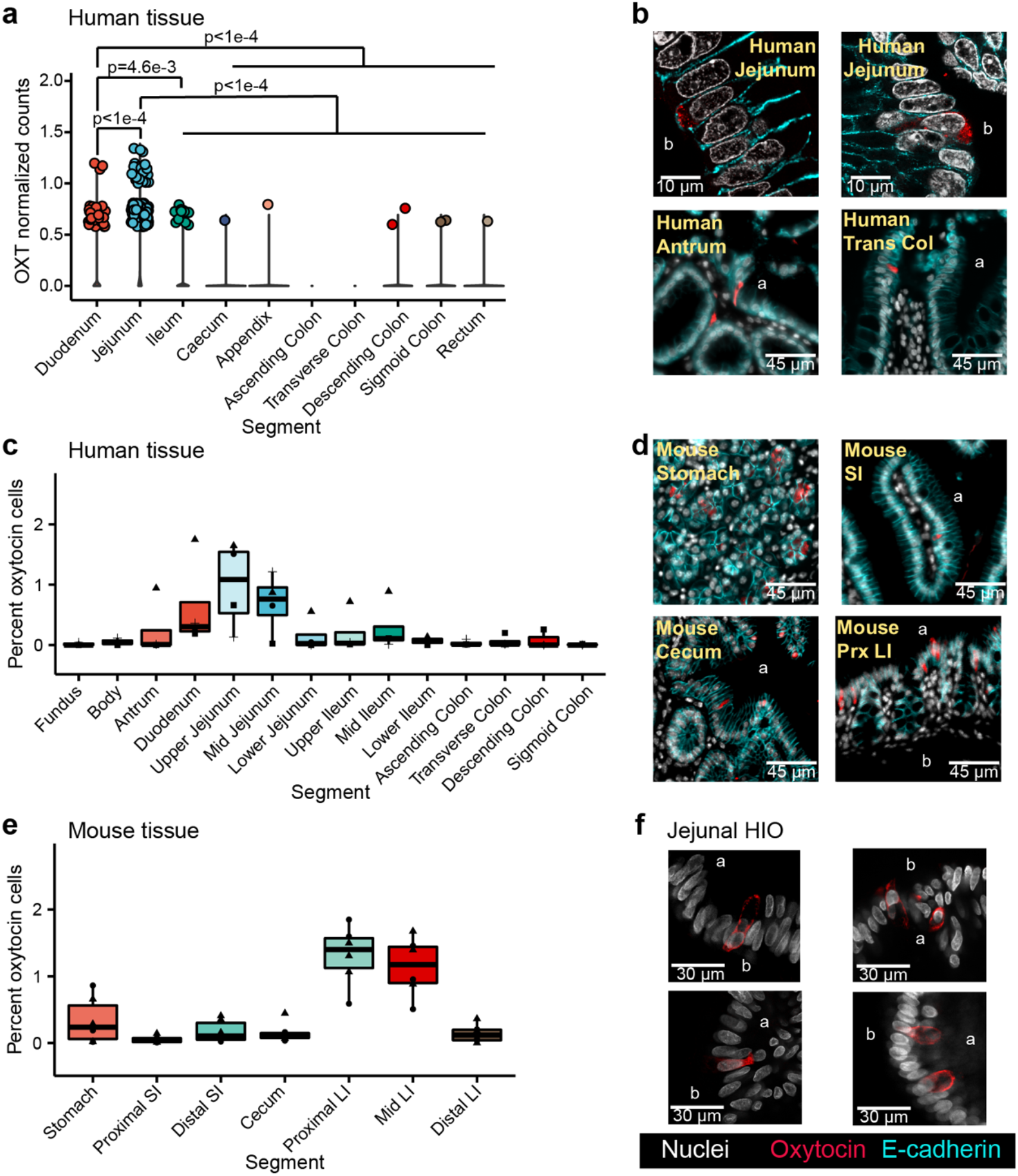
Oxytocin expression and production in the epithelium of the human and mouse gastrointestinal tract. **a)** Log normalized counts of oxytocin expression in various human intestinal epithelial tissues reported by the Gut Cell Atlas^35^. Significance values reflect the number of rarefactions (of 10,000) in which the comparison had a *p* value >0.05 by a Dunn Test with a Benjamini-Hochberg correction. These *p* values were similar whether the number of cells expressing oxytocin or oxytocin expression counts were used. Only *p* values <0.05 are shown. Oxytocin visualized by immunofluorescence imaging in **b)** 6 μm or 35 μm sectioned human jejunum, antrum, or transverse colon (Trans Col), **d)** mouse stomach, small intestine (SI), cecum, or proximal large intestine (Prx LI). Percentage of cells with oxytocin staining throughout the **c)** human or **e)** mouse intestinal tract. At least 3,000 nuclei were counted per segment. For c, shape denotes individual patient. For e, shape represents sex, male (triangle) or female (circle). **f)** Oxytocin visualized by immunofluorescence imaging in whole 3D J2-*NGN3* human intestinal organoids (HIO), differentiated but not induced for *NGN3*. In all images, DAPI stained nuclei are shown in white, oxytocin staining in red, and E-cadherin staining in cyan. Apical (a) and/or basolateral (b) sides are labeled. LI, large intestine. SI, small intestine. a: duodenum: *n* = 6,118 cells from 2 patients, jejunum: *n* = 2,791 cells from 4 patients, ileum: *n* = 5,439 cells from 4 patients, cecum: *n* = 16,334 cells from 6 patients, appendix: *n* = 4,648 cells from 4 patients, ascending colon: *n* = 6,051 cells from 4 patients, *n* = 6,437 cells from 6 patients for the transverse colon, descending colon: *n* = 1,712 cells from 3 patients, sigmoid colon: *n* = 7,480 cells from 7 patients for the, rectum: *n* = 16,405 cells from 3 patients; c: fundus, body, upper ileum, descending colon, and sigmoid colon: *n* = 3 unique patients, antrum, duodenum, mid jejunum, lower jejunum, mid ileum, lower ileum, ascending colon, and transverse colon: *n* = 4 unique patients; e: *n* = 6 mice per region.

To substantiate these findings, we performed immunostaining and fluorescence imaging on human intestinal tissue and observed oxytocin staining within the epithelial layer of the stomach, small, and large intestine (**Fig. 1b**), in both villi and crypts. In agreement with the human scRNA-Seq data, image analysis of at least 3,000 cells per intestinal segment indicated that the greatest number of oxytocin cells are present in the epithelium of the human upper small intestine (**Fig. 1c**). Oxytocin staining could be grouped into two general patterns: a somewhat granular signal within the cytoplasm as expected of hormones stored in vesicles, and a more diffuse signal that seemed to be between cells rather than around the nucleus. The latter staining pattern was observed more often in villi regions with dense oxytocin staining and could be indicative of staining within infiltrating immune cells (**Fig. 1c** versus **Extended Data Fig. 1b**). We also performed immunofluorescent staining on the mouse intestinal tract and observed oxytocin staining throughout the gut (**Fig. 1d**). However, in mice, we observed more oxytocin signal in the proximal and mid colon than in other regions of the intestine (**Fig. 1e**).

A few of the scRNA-Seq datasets we investigated were derived from intestinal organoids generated from adult gut epithelial stem cells. In the absence of other gut cell types, these stem cells differentiate to produce only the epithelial layer^43^. Therefore, if oxytocin is produced in these organoids, then the gut epithelium is sufficient to produce oxytocin independent of signals originating from the lamina propria, enteric neurons, or stromal layers. To test this hypothesis, we performed reverse transcription quantitative PCR (rt-qPCR) on human duodenal- and jejunal-derived organoids. In doing so, we observed *OXT* transcripts in differentiated (mature) organoids at higher levels than in undifferentiated (immature, mostly stem cells) organoid controls (**Extended Data Fig. 1c**). Similarly, upon imaging oxytocin in jejunal-derived organoids, we were able to readily observe staining in differentiated organoids but not undifferentiated organoids (**Extended Data Fig. 1f, Extended Data Fig. 1d, Extended Data Movie**). These data indicate that the intestinal epithelial layer is sufficient to produce oxytocin.

### *L. reuteri* promotes secretion of oxytocin from the intestinal epithelium

We next evaluated whether *L. reuteri* could stimulate secretion of oxytocin from the gut epithelium as *L. reuteri* is able to stimulate oxytocin in the hypothalamus^9^. To test this hypothesis, we applied *L. reuteri* cell-free conditioned medium, thereby comprising a mix of proteins, metabolites, nucleic acids and other molecules released by *L. reuteri* to mouse, pig, piglet, or human intestinal segments (**Fig. 2a**). In doing so, we observed significant secretion of oxytocin over medium alone controls throughout the human, pig, and piglet small intestine (**Fig. 2b-d, Extended Data Fig. 2a, b)** and in the mouse stomach and cecum (**Fig. 2e, Extended Data Fig. 2c, d**). Oxytocin secretion was observed in tissue isolated from both sexes in humans, mice, and piglets. For adult pigs, only females were tested. We next attempted to induce secretion of oxytocin from organoids in response to *L. reuteri*-conditioned medium (**Fig. 2a**). Given that oxytocin is a hormone, we performed this assay using an engineered jejunal human intestinal organoid line (J2-*NGN3* HIO) that can be induced to produce up to 40% enteroendocrine cells^44^. Indeed, we only observed a significant increase in oxytocin secretion driven by *L. reuteri*-conditioned medium versus control medium when using induced J2-*NGN3* HIOs (**Fig. 2f**). Notably, the ability of *L. reuteri* to stimulate oxytocin release was not shared by all bacteria, as *Bacillus subtilis* and *Escherichia coli* Nissle were unable to stimulate oxytocin release (**Extended Data Fig. 2e**).

**Fig. 2:**
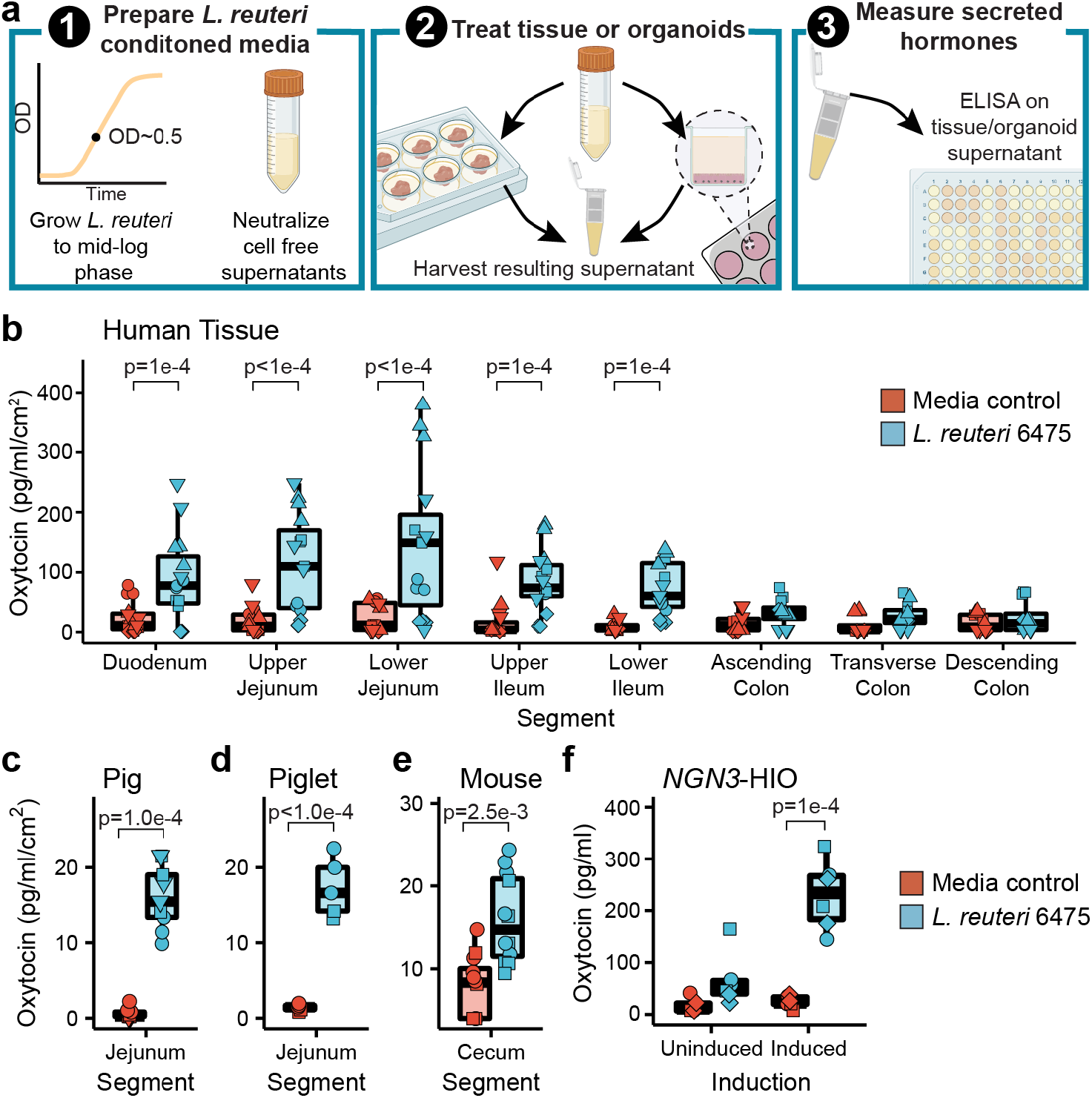
*L. reuteri*-conditioned medium promotes release of oxytocin from the gut epithelium. **a)** Workflow of secretion assays. *L. reuteri-*conditioned medium is prepared by growing *L. reuteri* to mid-log phase, spinning down the bacterial cultures, and harvesting the supernatant. The supernatant is then neutralized to ∼pH 7 and filter sterilized to remove cells while maintaining released products. For *ex vivo* tissue assays, *L. reuteri-*conditioned medium is placed onto intestinal tissue segments that have been washed free of luminal contents/fecal material. The tissue is then incubated with the *L. reuteri-*conditioned medium or *L. reuteri* growth medium control for 3 hours at 37ºC with 5% CO_2_. Afterwards, the resulting tissue supernatant is harvested, spun free of cells, and used in an ELISA. For organoid assays, organoids are prepared in a monolayer format and treated with *L. reuteri*-conditioned or growth medium control as for the *ex vivo* tissue. Figure made with BioRender. Oxytocin measured by ELISA and normalized by tissue surface area secreted from *ex vivo* **b)** human intestinal tissue **c)** pig jejunum, **d)** piglet jejunum, and **e)** mouse cecum. **f)** Oxytocin measured by Luminex secreted from uninduced and induced J2-*NGN3* HIOs. Point shape reflects unique patients (shown in triplicate) for human (b) pig (d), piglet (e), or organoid batch (in triplicate) (f). For mouse (e), points represent unique mice and shape denotes sex (females as circles and males as squares). Significance values were determined from the least squares means derived from linear or linear mixed models with pairwise comparisons corrected using a Benjamini-Hochberg multiple testing correction (see **Supplementary Tables 2** and **3**). b: *n* = 5 patients for the duodenum through lower ileum and *n* = 4 patients for the colon regions with three replicate tissues each (12 or 15 datapoints total); c: *n* = 3 animals per region and condition with three replicate tissues (9 datapoints total); d: *n* = 2 animals per region and condition with three replicate tissues (6 datapoints total); e: *n* = 12 animals per region and condition; f: *n* = 3 HIO batches with two replicate monolayers per condition (6 datapoints total).

### Oxytocin is produced by enterocytes in the intestinal epithelium

The reliance on an organoid model that has increased levels of enteroendocrine cells (EECs) suggested that oxytocin is made in an EEC, like other gut hormones^45^. However, we observed no increase in *OXT* transcripts, as measured by rt-qPCR, in induced J2-*NGN3* HIOs (**Fig. 3a**). In contrast, we observed a marked increase in *CHGA* transcripts encoding the neuroendocrine protein chromogranin A, which is known to be produced in EECs, in the induced J2-*NGN3* HIOs (**Fig. 3a**). Similarly, oxytocin positive cells were not increased by *NGN3* overexpression, as measured by flow cytometry, while chromogranin A positive cells were (**Fig. 3b**). Together, these results suggest that oxytocin is not produced by an EEC.

**Fig. 3:**
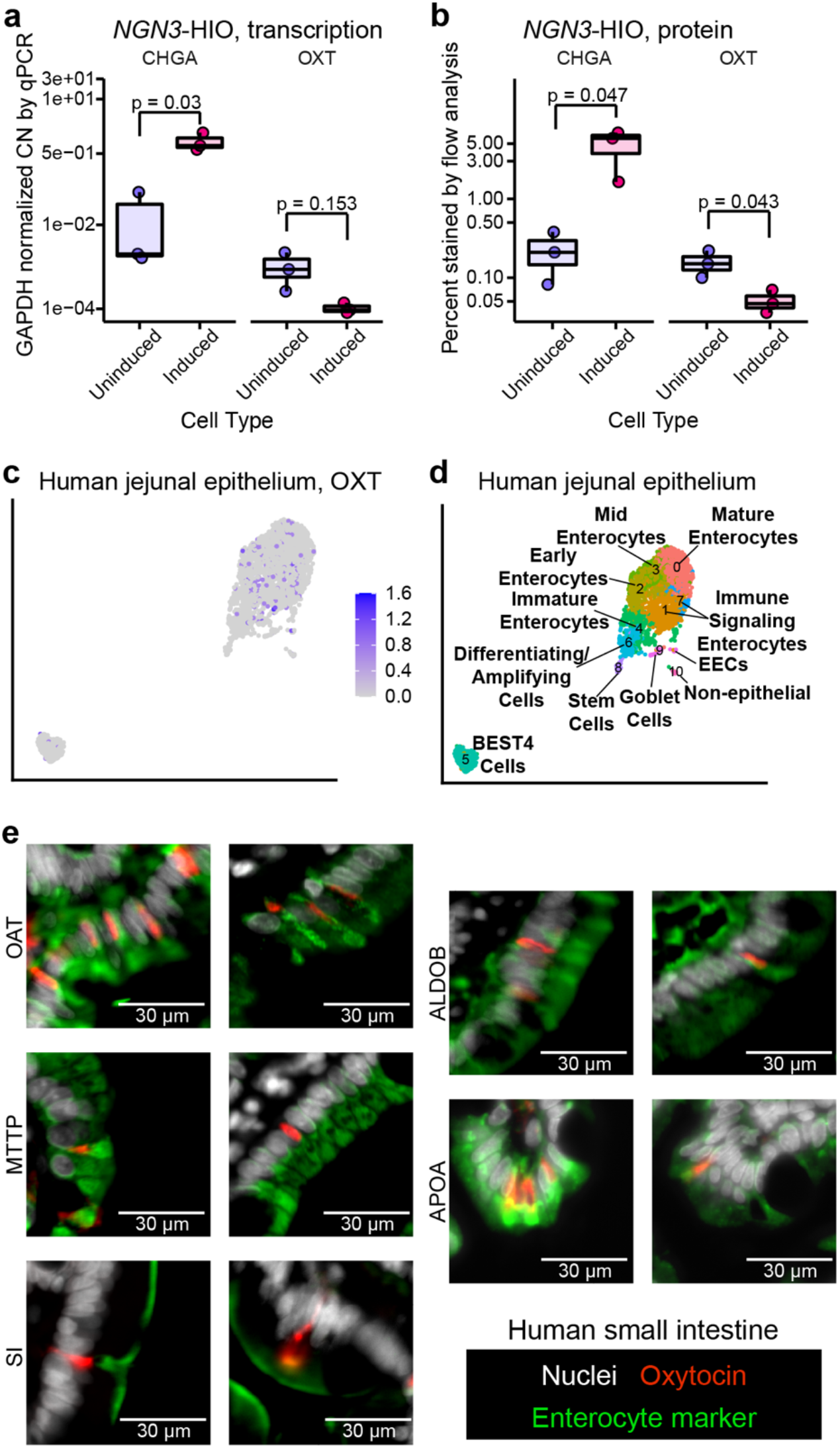
Oxytocin is an enterocytic hormone. **a)** Copy number (CN) of *CHGA* and *OXT* transcripts by qPCR and **b)** percent chromogranin A and oxytocin positive cells measured by flow cytometry in differentiated and induced J2-*NGN3* HIOs. For a, points represent averaged triplicate qPCR data each from a separate pooled batch of three 3D organoid wells, and where product could not be amplified, a *GAPDH* CN normalized value of 1e-5 was used. Only *p* values <0.05 are shown, which were determined from the least squares means derived from a linear model with pairwise comparisons corrected using a Benjamini-Hochberg multiple testing correction (see **Supplementary Tables 2** and **3**). UMAP of Gut Cell Atlas adult jejunal epithelium scRNA-Seq data^35^ labeled with **c)** oxytocin counts or **d)** identified cell clusters (see **Extended Data Fig. 3j** and **Supplementary Table 6**). **e)** Oxytocin (red) co-stained with enterocyte markers OAT, MTTP, SI, ALDOB, or APOA (green) in 6 μm sectioned human small intestinal tissue. Nuclei stained with DAPI are shown in white. a, b: *n* = 3 experiments; c, d: *n* = 2,791 cells from 4 different patients.

To determine which intestinal cell type produces oxytocin, we first turned to the EEC-enriched organoid scRNA-Seq dataset of Beumer and colleagues^46^. Surprisingly, *OXT* expression did not cluster with any of the major EEC subtypes (**Extended Data Fig. 3a, b, Supplementary Table 4**). Rather, *OXT* grouped alone in a cluster predominantly comprised of duodenal and ileal cells (**Extended Data Fig. 3c**) enriched in genes characteristic of enterocytes (*FABP2, FABP6, OAT, MTTP, RBP2, SI, APOA1, APOB*) (**Extended Data Fig. 3d, Supplementary Table 4**). In human small intestinal tissue, we confirmed that oxytocin does not co-localize with two markers of EECs: chromogranin A or neurotensin (**Extended Data Fig. 3e, f**). These results provide further evidence that oxytocin is not produced by an EEC but instead by an enterocyte.

To further test this hypothesis, we returned to the human tissue scRNA-Seq datasets of the Human Cell Landscape^39^ and Gut Cell Atlas^35^. In both datasets, we observed *OXT* expression predominantly within enterocyte clusters (**Fig. 3c, d, Extended Data Fig. 3g-j, Supplementary Tables 5, 6**). Furthermore, on co-staining human small intestinal tissue with oxytocin and enterocyte markers that span the entire villus, we observed co-localization of oxytocin and enterocyte-specific markers (**Fig. 3e**). Overall, these data suggest oxytocin is produced by a cell that functions as an enterocyte and we can classify this oxytocin as enterocytic.

To determine if other genes are uniquely expressed in oxytocin-producing cells, we compared the transcriptional profiles of oxytocin positive versus negative cells within the jejunum of the Gut Cell Atlas^35^ dataset. We observed that only *OXT* was significantly overexpressed (Wilcoxon rank sum test, log_2_ fold change = 1.1, padj < 1e-295), while *REG1A* and *OLFM4* (two stem cell markers) were significantly underexpressed (Wilcoxon rank sum test, log_2_ fold change < -1.1, padj < 0.05) in the oxytocin producing cells (**Supplementary Table 7**). Therefore, *OXT* expression alone appears to characterize cells that produce oxytocin.

### Secretin receptor signaling is necessary for oxytocin release by *L. reuteri* 6475

Having determined that oxytocin is not produced by an EEC, but that its secretion from organoids requires increased EECs, we next asked how *L. reuteri* stimulates the release of oxytocin. In the brain, hypothalamic oxytocin can be released by the peptide hormone secretin^47^, and both secretin and the secretin receptor are necessary for normal social behavior^47,48^. In the gut, secretin is produced in EECs by ‘S cells’^23^. Therefore, we hypothesized that gut produced secretin may promote oxytocin release. We explored this hypothesis by first testing if the addition of secretin would stimulate the release of oxytocin from intestinal organoids. Indeed, secretin applied to induced J2-*NGN3* organoids resulted in release of oxytocin at levels comparable to that of *L. reuteri*-conditioned medium (**Fig. 4a**). If secretin is sufficient to promote oxytocin release, we reasoned that we should be able to cause release of oxytocin from organoids not enriched in EECs by supplying exogenous secretin. We tested this hypothesis by applying secretin to non-engineered organoid lines derived from the human duodenum, jejunum, ileum, and ascending colon, derived each from a different individual. We observed secretion of oxytocin in each tested HIO line (**Fig. 4b**). Finally, we tested if application of secretin to whole intestinal tissue would promote oxytocin release. Consistent with organoid experiments, we observed release of oxytocin like that caused by *L. reuteri* (**Fig. 4c**).

**Fig. 4:**
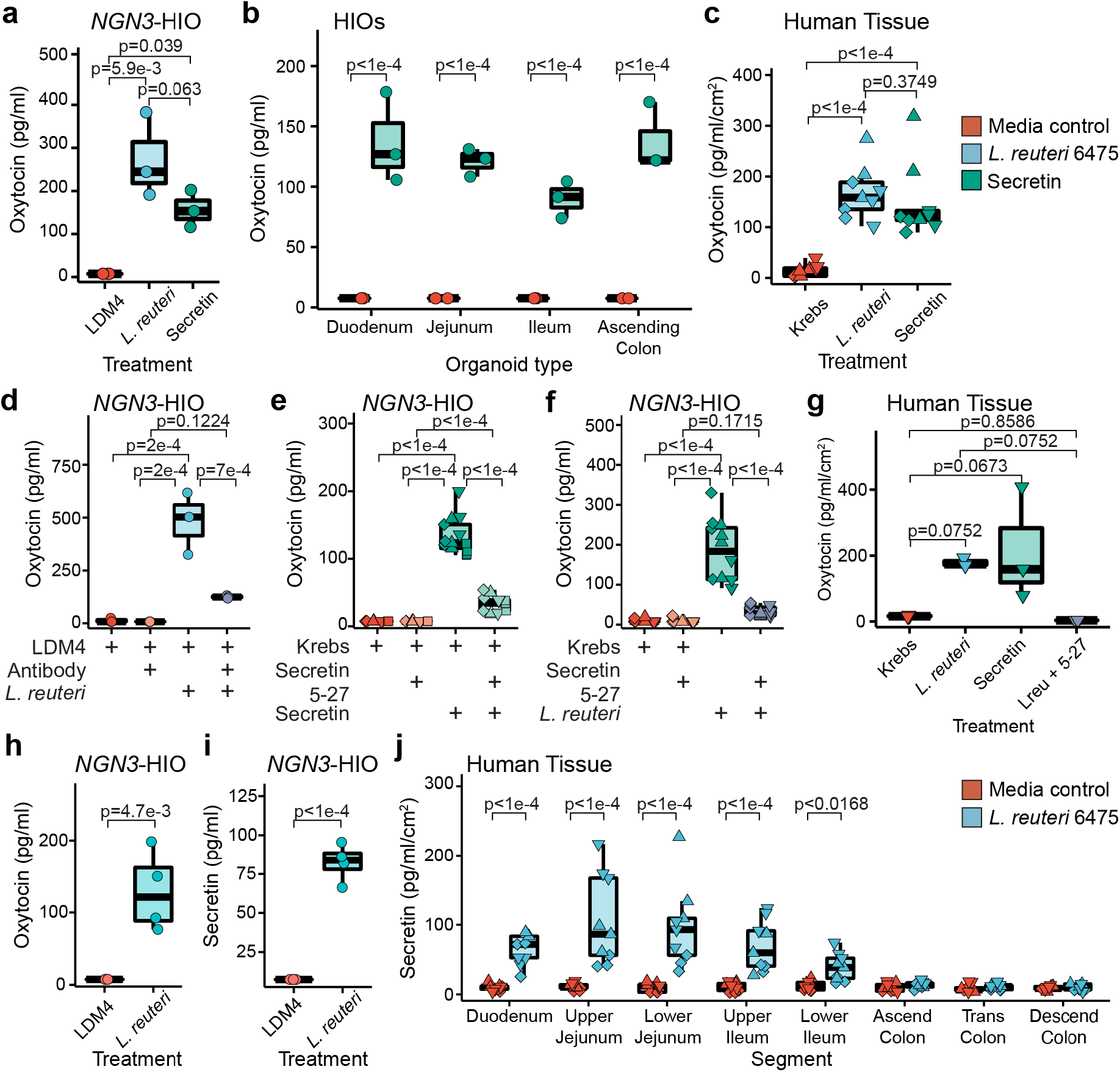
*L. reuteri* induces oxytocin secretion through release of secretin. Oxytocin measured by an ELISA released from **a)** induced J2-*NGN3* HIOs or **b)** non-engineered HIOs derived from different patients for different regions of the intestine treated with bacterial medium control (LDM4), *L. reuteri*-conditioned medium, or 1.2 ng/ml (120 pg total) of secretin. **c)** Oxytocin measured by an ELISA released from human mid-jejunal intestinal tissue treated with Krebs, *L. reuteri*-conditioned medium, or 2.5 ng/ml (12.5 ng total) of secretin. Oxytocin measured by an ELISA released from induced J2-*NGN3* HIOs treated with **d)** bacterial medium control (LDM4), secretin receptor antibody (1:100 dilution), and/or *L. reuteri*-conditioned medium. Oxytocin measured by an ELISA released from **e** and **f)** induced J2-*NGN3* HIOs or **g)** human mid-jejunal intestinal tissue treated with Krebs, 1.0 ng/ml (5 ng total) secretin 5-27, 2.5 ng/ml (12.5 ng total) of secretin, and/or *L. reuteri*-conditioned medium. **h)** Oxytocin or **i)** secretin measured by ELISA from the same induced J2-*NGN3* HIOs treated with bacterial medium control (LDM4) or *L. reuteri*-conditioned medium. **j)** Secretin measured by ELISA from human mid-jejunal intestinal tissue treated with bacterial medium control (LDM4) or *L. reuteri-*conditioned medium. Differential point shapes reflect multiple organoid batches (e, f) or unique patients (c, g, j). Significance values were determined from the least squares means derived from a linear or linear mixed model with pairwise comparisons corrected using a Benjamini-Hochberg multiple testing correction (see **Supplementary Tables 2** and **3**). a, b, d, h, and i: *n* = 3 replicate monolayers per condition; c and j: *n* = 3 patients with three replicate tissues per condition (9 datapoints total); e: *n* = 4 HIO batches with three replicate monolayers per condition (12 total datapoints); f: *n* = 3 HIO batches with three replicate monolayers per condition (9 total datapoints); g: *n* = 3 replicate tissues per condition from one patient.

To further verify that enterocytic oxytocin secretion is mediated through secretin, we blocked signaling through the secretin receptor in two ways. First, we blocked the secretin receptor on induced J2-*NGN3* organoids by pre-treatment with a blocking antibody prior to treatment with *L. reuteri*-conditioned medium. Inhibition of secretin receptor signaling significantly attenuated oxytocin release (**Fig. 4d**). Next, we tested whether a competitive inhibitor of the secretin receptor (secretin 5-27) would also block oxytocin release in induced J2-*NGN3* organoids. When secretin and secretin 5-27 were applied together, oxytocin release was abrogated (**Fig. 4e**). Similarly, when *L. reuteri-*conditioned medium was mixed with secretin 5-27, oxytocin was not secreted from induced J2-*NGN3* organoids (**Fig. 4f**). Finally, we treated human intestinal tissue with *L. reuteri-*conditioned medium supplemented with secretin 5-27 and observed loss of oxytocin secretion (**Fig. 4g**). Together, these findings demonstrate that signaling through the secretin receptor causes release of oxytocin from intestinal epithelial cells and suggests that *L. reuteri* 6475 induces oxytocin release via secretin.

### *L. reuteri* 6475-conditioned medium induces secretion of oxytocin and secretin

If *L. reuteri* 6475 is causing secretion of oxytocin via secretin, then *L. reuteri* 6475-conditioned medium should stimulate secretin release. To determine whether this was occurring, we measured both hormones from the same induced J2-*NGN3* HIO sample. In doing so, we observed co-release of secretin and oxytocin in response to *L. reuteri-*conditioned medium (**Fig. 4h, i**). We were also able to measure release of secretin from whole human tissue by *L. reuteri* (**Fig. 4j**) in the same intestinal regions in which we measured oxytocin (**Fig. 2b**). Collectively, these data show that *L*. reuteri-conditioned medium induces secretion of secretin (which is produced in EECs), and this secretin is necessary for *L. reuteri*-mediated release of oxytocin in the gut (**Fig. 5**).

**Fig. 5:**
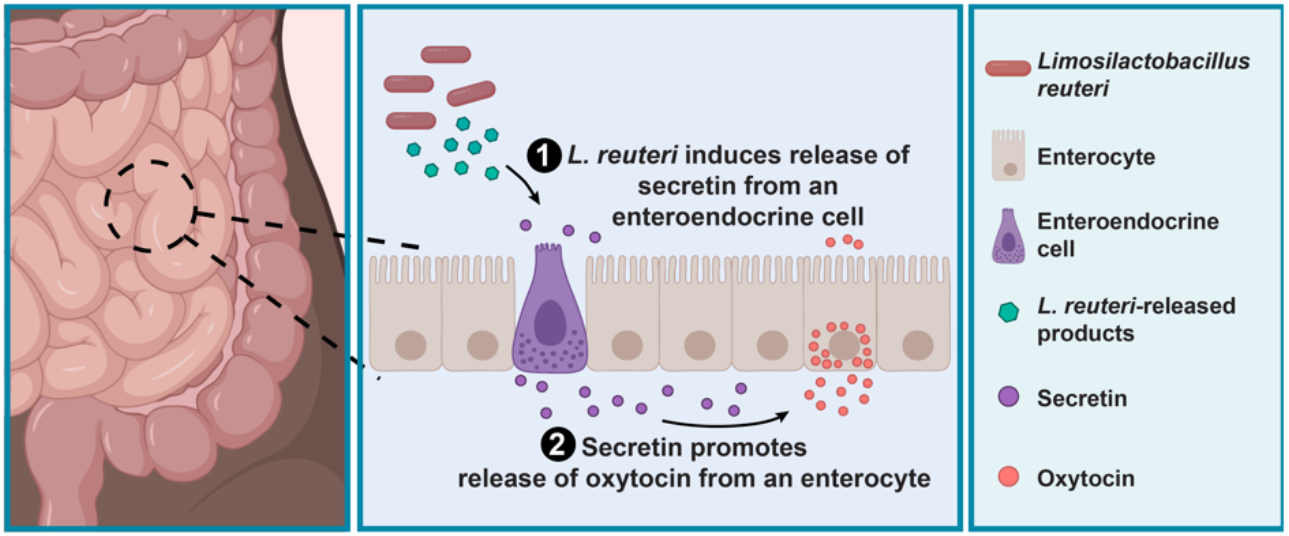
*L. reuteri* promotes the secretion of oxytocin from enterocytes through stimulation of secretin. Working model for *L. reuteri* mediated secretion of oxytocin. Oxytocin exists in enterocytes in the human small intestine. Its secretion is promoted by *L. reuteri* via secretin, which is produced in enteroendocrine cells. Figure made in BioRender.

## Discussion

The intestinal epithelium has evolved to rapidly sense and respond to beneficial and pathogenic microbes in the gut. How gut-resident microbes impact the function of other organs in the body and the specific role of the intestinal epithelium in these effects is an open area of research. The actions of the intestinal microbe *L. reuteri*, for example, result in beneficial effects on gut^1,30^, bone^11–17,29^, skin^5–7^, and brain physiology^8–10^. Signaling through the vagal nerve to the hypothalamus to release oxytocin in the brain is critical for the roles of *L. reuteri* in the brain^9^ and skin^5^, and oxytocin has similar effects on bone^21^, brain^10,49^, and the gut^19,20^ as *L. reuteri*. These observations suggest that oxytocin plays a critical role in many of *L. reuteri*’s systemic beneficial effects. Here, we report that oxytocin exists in the intestinal epithelium and its release is stimulated by *L. reuteri* via paracrine signaling by the hormone secretin.

Secretion of gut epithelial oxytocin can be stimulated by at least two factors in common with secretion of hypothalamic oxytocin: by *L. reuteri*^9^ and by secretin^47^. Secretin receptors are abundant in the intestine^50,51^ and may be present on an oxytocin producing cell or on a second cell type that coordinates oxytocin release. Interestingly, the link between oxytocin and secretin has been highlighted in several studies for the treatment of colitis^52^ and autism^53^. In mice, oxytocin was significantly secreted by *L. reuteri* in the stomach and cecum and variable among individuals in the large intestine. Secretin is predominantly produced in the proximal small intestine^54^, so oxytocin secretion in the mouse gut may be regulated by something other than secretin.

Oxytocin is recognized as an extraordinary hormone with a wide variety of beneficial effects^19–21,55–58^. Heretofore, these actions have been attributed almost exclusively to the release of oxytocin from the hypothalamus into the blood stream via the posterior pituitary gland or through activation of other regions in the brain^31^. However, oxytocin is also observed in a variety of other tissues including the thymus^59^, heart^60^, uterus^61^, placenta^61^, testes^62^, and intestinal neurons^32^. The roles of these locally produced oxytocin are not well understood. In our investigation of human and mouse intestinal tissue, we discovered that oxytocin is also produced within the intestinal epithelium in both villi and crypts.

Surprisingly, oxytocin was not produced in an enteroendocrine cell but rather in a subset of enterocytes. Oxytocin is not the only hormone known to be produced from an enterocyte: FGF19 (mouse Fgf15) is also secreted from an enterocyte cell type^63,64^. Oxytocin was the only gene distinguishing the population of oxytocin-producing cells from other enterocytes. Interestingly, in the hypothalamus, oxytocin is also the only gene enriched in oxytocin producing neurons^65^. The presence of oxytocin in enterocytes suggests a potential function relating to food sensing or metabolism. Indeed, some of the functions attributed to oxytocin are in glucose and lipid metabolism^66^.

Our work reveals that the distribution of oxytocin in the gut differs between humans and mice: humans had much more in the upper small intestine, whereas mice had much more in the upper large intestine. Disparities in hormone density along the gastrointestinal tract between mice and humans has been noted for other gut hormones including PYY and GLP-2^67^. What causes these specificities remains to be understood. In humans the enrichment of oxytocin in the upper small intestine suggests it could be regulated by diet, bile, small intestinal hormones like secretin^68^, and small intestinal microbes including *L. reuteri*^33^.

*L. reuteri*-mediated release of oxytocin, we demonstrate, is facilitated by secretin signaling. The mechanism through which *L. reuteri* induces release of secretin is currently unknown. *L. reuteri* is known to cause secretion of histamine^2^, which can stimulate oxytocin secretion^69,70^ in the hypothalamus. Previously, it was reported that *L. reuteri* promotes synthesis of tetrahydrobiopterin^10^, which promotes social behavior with dependency on the oxytocin receptor^10^ and mediates oxytocin release in the brain^71,72^. Whether there is a relationship between intestinal epithelial oxytocin and tetrahydrobiopterin remains to be determined.

Our findings raise major outstanding questions regarding enterocytic oxytocin. First, is there a relationship between enterocytic oxytocin and hypothalamic oxytocin secretion? Does release of oxytocin from the intestinal epithelium enhance secretion of oxytocin from the hypothalamus? Alternatively, does *L. reuteri*-mediated release of secretin directly or indirectly promote oxytocin release in the brain, similar to how circulating cholecystokinin can activate brain oxytocin^73–76^? Second, what is/are the functions of enterocytic oxytocin? One possibility is that enterocytic oxytocin acts locally in the gut. Whole body oxytocin knockout mice have increased gut motility, increased intestinal inflammation, less proliferative epithelial stem cells, and increased epithelial permeability^19^. Another possible function of enterocytic oxytocin is that it has a function related to food metabolism. Whether any of the other roles of total body oxytocin are driven by enterocytic oxytocin remains unknown.

Altogether, this work demonstrates that oxytocin is produced and secreted from enterocytes in the intestinal epithelium. This discovery identifies a new mechanism by which gut microbes can regulate host physiology within the gut. Further work is needed to define this host-microbe interaction on the molecular level and consequences of this interaction.

## Methods

### scRNA-Seq analysis

Unless noted otherwise, we determined the number of oxytocin cells present in published scRNA-Seq datasets by enumerating the number of cells reported as expressing oxytocin without filtering the data.

For more detailed analysis, data from Beumer et al^46^ were loaded into the Seurat package in R (v 3.2.1)^77^. Cells were filtered so to only include those with greater than 1,100 unique transcripts, and less than 9,000 unique transcripts, similar to methods used to process the data presented in the original publication^46^. Data were log normalized using the NormalizeData function, variable features found using 2,000 features and the “vst” method, and data were scaled using default settings for the ScaleData function. Using the ElbowPlot function, we selected 17 dimensions to reduce and cluster the data using the FindNeighbors function. Clusters were found using the FindClusters function with resolution = 0.9 and data were plotted using a UMAP reduction. Informative plots were made using the FeaturePlot and VlnPlot functions. Marker genes for each cluster were found using the FindAllMarkers function with min.pct = 0.25, logfc.threshold = 0.25, only.pos = TRUE, and only genes with an adjusted p-value (from a Bonferroni correction) of <0.05 and a log fold change greater 1 or less than -1 were further considered.

For the scRNA-Seq data of the Human Cell Landscape^39^, we used background gene expression corrected data as provided. Data were similarly processed as for the Beumer et al 2020 data, such that we filtered these data to only include cells with greater than 50 transcripts, greater than 70 but less than 1,800 unique transcripts, and less than 45% mitochondrial transcripts. We used 5,000 features and varied the number of dimensions according to the output from the ElbowPlot function in Seurat and the resolution so to maximize the number of clusters while keeping the maximum modularity greater than 0.75. From this analysis, we identified cells belonging to the epithelial layer by first identifying markers for each cluster using the FindAllMarkers in Seurat and cross-checking these markers against the annotations presented in the Human Cell Landscape online tool (https://db.cngb.org/HCL/). The epithelial cells were then processed using the same pipeline to produce a UMAP, adjusting the number of dimensions and resolution as necessary. To compare oxytocin expression among intestinal segments, data were SCTransformed with the options method = “glmGamPoi”, vars.to.regress = percent mitochondrial reads, return.only.var.genes = FALSE, and min_cells = 1.

For the adult epithelial scRNA-Seq data from the Gut Cell Atlas^35^, we filtered the data to exclude batches containing less than 50 cells. The data were split into batches and then transformed with SCTransform as for the Human Cell Landscape data. This SCTransform data were carried through the remainder of the pipeline to produce a UMAP.

To compare the expression of oxytocin across intestinal sites, we rarefied the SCTransformed data for oxytocin in the Human Cell Landscape dataset to 1281 cells, thereby excluding the ascending colon and epityphlon, and the Gut Cell Atlas data to 1712 cells. For each dataset, we performed the rarefaction 10,000 times. Each iteration, expression difference significances among intestinal regions were determined by a Kruskal-Wallis test and a Dunn test was used for pairwise comparisons with a Benjamini-Hochberg multiple testing correction. Reported *p* values are the fraction of iterations in which the given comparison had a *p*>0.05 from the Dunn test.

To find other genes that may be enriched in oxytocin positive cells, we used the FindMarkers function in Seurat using a Wilcoxon rank sum test on oxytocin positive and negative cells in the Gut Cell Atlas jejunum dataset after batch normalization and integration. Genes with log_2_ fold change ≥1 or ≤-1 and with an adjusted *p* <0.05 were considered as significant.

### Ex vivo *tissue*

#### Human

Intestines were acquired through the organ donation group LifeGift (Houston, TX, USA). Whole intestines were delivered on ice within 1 hour of removal. Intestinal regions were excised (sizes between 1.4 to 8 cm^2^), washed in cold PBS, and used for secretion assays or imaging (see below).

#### Pig

Intestines from pregnant sows and piglets were a gift of Douglas Burrin and Barbara Stoll, which were collected in accordance with IACUC policies. Collected intestines were treated as were for human tissue. Pieces ∼ 7.07 cm^2^ from pigs and 2.5 to 6 cm^2^ from piglets in size were used for secretion assays. Piglets used were one male and one female with ages 3 and 4 weeks respectively. Sows were within a few days from natural delivery.

#### Mouse

Mouse tissue was acquired from conventional mice both housed in a specific pathogen free facility on standard chow diet under Baylor College of Medicine protocol AN:8471. Male and female adult (age matched, 9 to 12 weeks old) mice were euthanized by CO_2_ affixation followed by cervical dislocation. Whole stomach, small intestine, cecum, and large intestine were removed and washed with PBS.

### Tissue immunofluorescence

Tissue regions after excision and washing were placed in TRU-FLOW tissue cassettes (Fisherbrand, Sugar Land, TX, USA) in 4% paraformaldehyde in PBS, with a volume at least 50x that of the tissue, at 4°C. After 18 to 24 hours, cassettes were rinsed twice in PBS and transferred to 70% ethanol. Tissue was embedded in paraffin and sectioned to 6 μm or 35 μm sections by the Digestive Diseases Center Cellular and Molecular Core at Texas Children’s Hospital. Sections were deparaffinized in three changes xylene for 10, 10, and 15 mins and hydrated through two 10 min changes in each of 100%, 95%, 70% and 50% ethanol followed by two 10 min washes in diH_2_O. Antigen retrieval was performed using a 0.01M sodium citrate buffer, pH 6.0 in the Retriever 2100 (Electron Microscopy Sciences, Hatfield, PA, USA) with resting in the buffer overnight to cool to room temperature. Slides were rinsed three times in diH_2_O, with the last rinse lasting 15 mins. Pap pen (Sigma-Aldrich, St. Louis, MO, USA) isolated areas on the slides were then placed in PBS for 3 minutes before permeabilizing with 0.5% Triton-X 100 (Sigma-Aldrich, St. Louis, MO, USA) for 20 mins. Slide regions were washed in PBS for 5 mins and then blocked with 10% normal goat (Jackson ImmunoResearch, West Grove, PA, USA) or donkey serum (Sigma-Aldrich, St. Louis, MO, USA) in PBS-0.05% Tween 20 (PBS-T, Genesee Scientific, El Cajon, CA, USA) for 1 to 2 hours. Next primary antibodies made in 1% normal goat or donkey serum in PBS-T were added to the slide regions, and the slides were incubated at 4°C in a humid slide staining tray (Newcomer Supply, Middleton, WI, USA) overnight. Primary antibodies were washed three times for 10 mins each in PBS-T.

Secondary antibodies with NucBlue Fixed Cell ReadyProbes (Invitrogen/Life Technologies, Carlsbad, CA, USA) and Alexa-647 conjugated E-cadherin (BD Biosciences, Franklin Lakes, NJ, USA) were applied diluted in 1% normal goat or donkey serum in PBS-T for 1 hour at room temperature and slides were kept dark within the humid slide staining tray. If a mouse raised primary antibody was used, then secondary antibodies were applied alone. In this case, secondary antibodies were washed three times for 10 mins each in PBS-T and then the slide regions were blocked in 5% normal mouse serum (Jackson ImmunoResearch, West Grove, PA, USA) in PBS-T for 1 hour after which NucBlue Fixed Cell ReadyProbes and Alexa 647 conjugated E-cadherin were applied diluted in PBS-T for 1 hour at room temperature with slides kept dark within the humid slide staining tray. After the last antibody/probe incubation, slides were washed three times for 10 mins each in PBS. Slides with 6 μm sections were mounted in ProLong Glass Antifade Mountant (Thermo Fisher Scientific, Sugar Land, TX, USA) and those with 35 μm sections in SlowFade Glass Soft-set Antifade Mountant (Thermo Fisher Scientific, Sugar Land, TX, USA). Primary and secondary antibodies and their dilutions are listed in **Supplementary Table 8**. Slides were imaged on either a Revolve (Echo, San Diego, CA, USA) fluorescence microscope, Axio Observer (Zeiss, Pleasanton, CA, USA) fluorescence microscope, or LSM 880 with Airyscan (Zeiss, Pleasanton, CA, USA). Image processing was accomplished in Zen Blue (Zeiss, Pleasanton, CA, USA) and/or Fiji^78^.

### Image quantification of oxytocin staining

For quantification at least 40 fields of view with 20% overlap were imaged on a Zeiss Axio Observer at 20x magnification. The images were stitched using the Zen Blue software, allowing for 5 to 20% overlap of the fields of view. Oxytocin staining in the stitched images were manually counted. Nuclei were counted with a custom script in MatLab. Briefly, first, E-cadherin staining (far red channel) was used to create a mask of the epithelial layer. Then a mask of the nuclei was created from the DAPI staining (blue channel). Then the E-cadherin mask and nuclei masks were overlapped to identify nuclei within the epithelial layer. Finally, the number of nuclei was estimated by dividing a pre-estimated average nuclei area by the total nuclei area in the epithelium. Finally, percent oxytocin cells were calculated as the percent of oxytocin positive cells of the total epithelial nuclei.

### Organoid culture and maintenance

Organoids (duodenal D103, jejunal J11 and J2-*NGN3*, ileal IL104, and ascending colon ASC209) were acquired from the Digestive Diseases Center Organoid Core at Baylor College of Medicine. Organoids enriched in enteroendocrine cells (J2-*NGN3*) were previously generated and characterized^44^. All other organoids are non-engineered organoid lines each from a different patient as indicated by numeric designation in the line name. Briefly, organoids were embedded in Matrigel (Corning, NY, USA) in a 24 well plate (Nucleon Delta, Thermo Scientific, USA) and cultured in complete medium with growth factors (CMGF+) supplemented with 50% (v/v) Wnt3a conditioned medium, 10 µmol Y-27632 Rock inhibitor and 100 µg/ml normocin (InvivoGen, San Diego, USA) medium in a humidified 5% CO_2_ incubator at 37ºC. J2-*NGN3* organoids were also supplemented with 200 µg/ml Geneticin (Gibco, Grand Island, NY). Organoids were routinely passaged every ∼6 to 10 days. All organoid cultures were passaged less than 60 times prior to downstream experimental manipulations.

### 3D imaging of organoids

Organoids were seeded in 3D in Matrigel (Corning) plugs as for passaging organoids. Following seeding, organoids were given hW-CMGF+ medium^44^ supplemented with 10 µmol Y-27632 Rock inhibitor, and for J2-*NGN3* organoids, 200 µg/ml Geneticin for 2 days. After two days, media were changed to according to organoid treatment group: undifferentiated: hW-CMGF+ medium, 10 µmol Y-27632 Rock inhibitor, and for J2-*NGN3* organoids 200 µg/ml Geneticin; differentiated and uninduced: differentiation medium^44^ with 10 µmol Y-27632 Rock inhibitor; differentiated and induced (J2-*NGN3* organoids only): differentiation medium, 10 µmol Y-27632 Rock inhibitor, and 1.0 µg/ml doxycycline (as doxycycline monohydrate, Sigma-Aldrich, St. Louis, MO, USA, dissolved in DMSO). These media conditions (at 500 µl) were continued for 4 days with daily media changes. After 4 days, 3D organoids were prepared for imaging^79^. Epithelial cell boundaries were imaged by staining with Alexa 647 conjugated E-cadherin, and nuclei were stained with 0.07x NucBlue Fixed Cell Stain ReadyProbes (Invitrogen, USA) with secondary antibody application. Primary and secondary antibodies and their dilutions are listed in **Supplementary Table 8**.

### Bacterial culture and preparation of conditioned medium

*Limosilactobacillus reuteri* 6475 (ATCC PTA 6475) was cultured overnight (∼16 hours) at 37ºC in deMan, Rogosa, Sharpe broth (MRS; BD Difco, Franklin Lakes, NY, USA). *Escherichia coli* Nissle and *Bacillus subtilis* were cultured overnight at 37ºC in Luria-Bertani broth (BD Difco, Franklin Lakes, NY, USA). To prepare conditioned media, strains were subcultured in LDM4, a fully defined medium^80^ with a starting optical density of 0.1 and cultured at 37ºC until an optical density of 0.5. Cells were collected by centrifugation and the supernatant was neutralized with 10M sodium hydroxide to pH ∼7.0 and filtered through a 0.22 µM PVDF filter (MilliporeSigma, Burlington, MA, USA).

### rt-qPCR of organoids

Organoids were plated in 3D in Matrigel and cultured for one day in hW-CMGF+ medium supplemented with 10 µmol Y-27632 Rock inhibitor, 100 µg/ml normocin, and for J2-*NGN3* organoids, 200 µg/ml Geneticin. Following, organoids were maintained in hW-CMGF+ medium both supplemented with 200 µg/ml Geneticin for J2-*NGN3* HIOs, switched to differentiation medium^44^ supplemented with 200 µg/ml Geneticin for J2-*NGN3* HIOs, or switched to differentiation medium^44^ supplemented with 1 µg/ml doxycycline for *NGN3* induction. Per biological replicate, three replicate wells per media were used. Media were changed daily for two additional days. To harvest cells for RNA extraction, replicate organoid wells were combined in ice cold dPBS and spun at 200x*g* for 5 mins at 4°C. Following removal of the supernatant containing dPBS and Matrigel, 1 ml TRIzol (Invitrogen, Carlsbad, CA, USA) was added and cells were vortexed for 30 secs at max speed and incubated at room temperature for 5 mins. Extraction continued with the addition of 200 µl chloroform, vortexing for 20 secs at max speed and with incubation at room temperature for 3 mins. To separate the phases, cells were spun at 14,000x*g* for 15 mins at 4°C. The aqueous (upper) phase was moved to a fresh tube and mixed with an equal volume of 70% ethanol. Samples were transferred to spin columns from the RNeasy Isolation Kit (Qiagen, Hilden, Germany), and RNA was purified using the manufacturer’s instructions with elution in 30 µl RNase free water. RNA was quantified and purity was determined using a DeNovix DS-11 Spectrophotometer (DeNovix, Wilmington, DE, USA). DNA was removed by treating 200 ng of RNA with TURBO DNase (Invitrogen, Carlsbad, CA, USA) according to the manufacturer’s instructions. cDNA was made from 20 ng of the resulting RNA using SuperScript III (Invitrogen, Carlsbad, CA, USA) per the manufacturer’s instructions with 50 ng random hexamers. Reactions were done in triplicate and combined prior to qPCR. No reverse-transcriptase controls were prepared as well.

Quantitative PCR reactions were performed in triplicate and consisted of 1 µl cDNA from above, 0.25 µl 20 mmol/L forward and reverse primers, 10 µl PowerUp SYBR Green Master Mix (Applied Biosystems, Foster City, CA, USA), and 8.5 µl nuclease free water. Reactions were performed on a QuantStudio 3 real-time thermocycler (Applied Biosciences, Foster City, CA, USA), and conditions consisted of 50°C for 2 mins, 95°C for 10 mins, 40 cycles of 95°C for 15 secs and 60°C for 1 min, after which fluorescence was measured. Melt curve analysis was performed by heating between 95°C and 60°C. No DNA and no reverse transcriptase controls were included. Cycle threshold values were determined using the QuantStudio software. Data analyses were performed in R version 3.5.2^81^. Briefly, cycle threshold values from triplicate qPCRs were averaged for a biological replicate. Undetermined cycle threshold values (target undetected within 40 cycles) were removed, and only averages resulting from replication of two or greater reactions were carried further. Cycle threshold values were normalized against that for *GAPDH* and converted to copy number: 2^(Ct.GAPDH-Ct.Target). Plotted data are replicate biological experiments (n≥ 2) and were graphed. Significance of expression organoid groups per organoid line were computed using linear models with a Benjamini-Hochberg multiple comparison adjustment using the lm function in R^81^. Least-squares means estimates were computed with the emmeans function in the emmeans package^82^ using a Benjamini-Hochberg multiple testing correction (see **Supplementary Tables 2** and **3**). Primer sequences are provided in **Supplementary Table 9**.

### Tissue secretion assays

For mouse secretion assays, the entire stomach, small intestine, cecum, and large intestine were removed, flushed/washed free of contents with PBS using a blunt 18-gauge needle (small and large intestine) or by filleting the tissue (stomach and cecum). The small and large intestine were then filleted. Each intestinal segment was then placed into 1 ml of treatment in a 12 well plate. Plates were incubated at 3 hours at 37ºC in a humidified 5% CO_2_ cell culture incubator.

Supernatants were collected, spun down at 1,000xg, and frozen at -20ºC or used directly in an ELISA. For human or pig tissue, tissue was placed into 5 ml of treatment in a 6 well plate and subsequently handled as for the mouse tissue.

### Organoid secretion assay

Organoid monolayers were produced by coating 96 well plates in 25 µl/ml Matrigel and seeding them with two 3D wells of J2-*NGN3* organoids per one 96 well. After two days recovery in CMGF+, monolayers were differentiated using differentiation medium^44^ supplemented with Rock inhibitor and 1 µg/ml doxycycline. After 5 days differentiation, media were removed and replaced with treatment. Monolayers were incubated at a humidified 5% CO_2_ incubator at 37ºC for 3 hours. Following, supernatants were removed and frozen at -20ºC.

### Quantitation of secreted hormones

Oxytocin in the tissue or organoid supernatants were measured using an ELISA (Enzo Life Sciences, USA, ADI-900-153A-0001) or Luminex panel (Millipore Sigma, USA, HNPMAG-35K). Secretin was similarly measured using an ELISA (Biomatik, USA, EKU07226). For pig tissue, supernatants were concentrated 5x for the ELISA by lyophilizing. Extrapolated values from the assays less than the limit of detection were given the value of half the lowest standard in the standard curve for the ELISA. Data were analyzed using either a linear or a linear mixed model. Linear mixed models were used for data derived from human or pig intestinal tissue with the patient or pig variable added as a random effect, for the piglet data, sex was added as a random effect as there was only a single male and a single female piglet, and for organoids where multiple organoid batches were included as a random effect. For mouse data, other organoid data and for those with multiple organoid lines, linear models were used with organoid line included as a variable. The specific models used are provided in **Supplementary Table 2**. Linear mixed models were run using the lmer function in lme4 package^83^ with REML = FALSE and the control optimizer = “bobyqa” and linear models with the lm function in R^81^. Least-squares means estimates were computed with the emmeans function in the emmeans package^82^ using a Benjamini-Hochberg multiple testing correction (see **Supplementary Table 3**).

### Quantitation of hormones by flow cytometry

Organoids in 3D Matrigel plugs were washed once with cold dPBS, then incubated in Cell Recovery Solution (Corning, Corning, NY, USA) at 4°C on a horizontal shaking platform for 30 to 60 minutes after mechanical disruption of the matrigel plugs. Following, the organoids were collected into a centrifuge tube and spun down at 200xg at 4°C for 5 minutes. Organoids were resuspended in 0.05% trypsin (diluted in dPBS from 0.25% trypsin-EDTA, phenol red, Gibco, USA), using 1 ml for 9 to 12 wells from the original 24 well plate housing the 3D Matrigel plugs, and incubated at 37°C for 10 mins. Trypsin was inactivated by adding 2x volume of cold CMGF-with 10% FBS. Organoids were fully dissociated by pipetting 100 times with a 1 ml pipette tip. Following, the organoid solution was filtered through a 40 µm strainer and centrifuged at 400xg for 5 minutes. Cells were resuspended in 2 ml CMGF- and spun again at 400xg for 5 minutes. Cells were resuspended in 0.25 ml of 4% paraformaldehyde per staining condition, transferred to a 5 ml round bottom polypropylene tube, and incubated at 4°C for 45 mins to fix the cells. Cells were then permeabilized by directly adding 16x volume (4 ml) of 0.1% Tween 20 in dPBS and incubating at 4°C for 10 mins. Cells were spun as before, resuspended in 1 ml 0.1% Tween 20 in dPBS and stored at 4°C for 1 to 2 days or the protocol was immediately continued.

To block cells prior to staining, cells were spun at 400xg for 5 minutes and resuspended in 1 ml per staining condition of 0.1% Triton-X 100, 5% donkey serum, 1% BSA in dPBS and incubated at 4°C for 20 minutes. For primary antibody staining, cells were spun as before, resuspended in 0.25 ml primary antibodies diluted in 0.1% Triton-X 100, 1% BSA in dPBS, and incubated at 4°C overnight. Following the cells were washed by directly adding 3 ml of 0.1% Triton-X 100, 1% BSA in dPBS incubating at 4°C for 3 minutes, and spinning down the cells as before. Cells were washed an additional 3 times in a similar manner, waiting 10 to 30 minutes between washes. For secondary antibody and DAPI staining, cells were spun as before, resuspended in 0.25 ml secondary antibodies and DAPI diluted in 0.1% Triton-X 100, 1% BSA in dPBS, and incubated at 4°C overnight in the dark. Cells were washed as before and resuspended in 400 µl 2% BSA, 2 mM EDTA, 2 mM sodium azide in dPBS and stored at 4°C in the dark until flow cytometry analysis.

Flow analysis was completed on an Attune NxT (Invitrogen, USA). Data were analyzed using FlowJO (v10, BD Biosciences, USA). Single cells were identified using 1) forward scatter area VS side scatter area, 2) forward scatter area VS forward scatter height, 3) side scatter area VS side scatter height, 4) forward scatter area VS violet channel area (for DAPI). Cells representing those in G1 were selected and carried forward. 100,000 to 500,000 DAPI+ single G1 cells were analyzed per sample and at least 60,000 per staining control. OXT+ and CHGA+ cells were identified by gating on the blue channel (for OXT) or yellow channel (for CHGA) such that the percent of cells in samples lacking the primary antibody for OXT and CHGA was less than 0.05%. See **Extended Data Fig. 4** for an example of the gating strategy.

### Statistical analyses

All data analyses were conducted in R (v 4.1.2)^81^. All boxplots show the median and the lower and upper quartiles. All data points are shown. All tailed tests are two-tailed. See specific method sections and **Supplementary Tables 2** and **3** for details. Significance values of linear mixed models were derived by performing an ANOVA of the model and a null model lacking all fixed effects. Exact *n* values are provided in the figure legends.

### Data availability

Data and R and MatLab scripts used to generate the figures are available at https://github.com/sdirienzi/Lreuteri_OXT.

## Acknowledgments

We acknowledge Javier Nieto, Mary Estes, Sarah Blutt, Shelly Buffington, Carolyn Bomidi, Victoria Poplaski, Colleen Brand, Erika Nachman, Micah Forshee, Madeline Bresson, Joseph Hyser, Douglas Burrin, Barbara Stoll, Caitlin Vonderohe, Pamela Parsons, Susan Venerable, Xi-Lei Zeng, Xiaomin Yu, Brian McDaniel, Nick Adair, and Mike Dehart for assistance and intellectual support. This project was supported by BioGaia (RAB), the Weston Foundation (RAB), and NIH grants P30 DK056388 (SCD and HAD), T15LM007093 (SCD), and F32 AI136404 (HAD). This project was supported in part by NIH grant DK056338 (Cellular and Molecular Core, Functional Genomics and Microbiome Core, and Gastrointestinal Experimental Model Systems Core), which supports the Texas Medical Center Digestive Diseases Center. This project was supported by the Cytometry and Cell Sorting Core at Baylor College of Medicine with funding from the CPRIT Core Facility Support Award (CPRIT-RP180672), the NIH (P30 CA125123 and S10 RR024574). This project was supported by the Optical Imaging and Vital Microscopy Core at Baylor College of Medicine.

## Author contributions

HAD, RAB, SCD conceptualized, designed the experiments, and interpreted the data. HAD, JL, AT, SCD performed the experiments and analyzed the data. SCD prepared the original draft. All authors reviewed the draft. HAD, JL, RAB, SCD revised the draft. HAD, RAB, and SCD acquired funding and materials for the project.

## Competing interests

Funding was provided by BioGaia (RAB), with whom we have a provisional patent (HAD, RAB, SCD).

## Materials & Correspondence

Requests should be directed to RAB and/or SCD.

---

**Extended Data Fig. 1:**
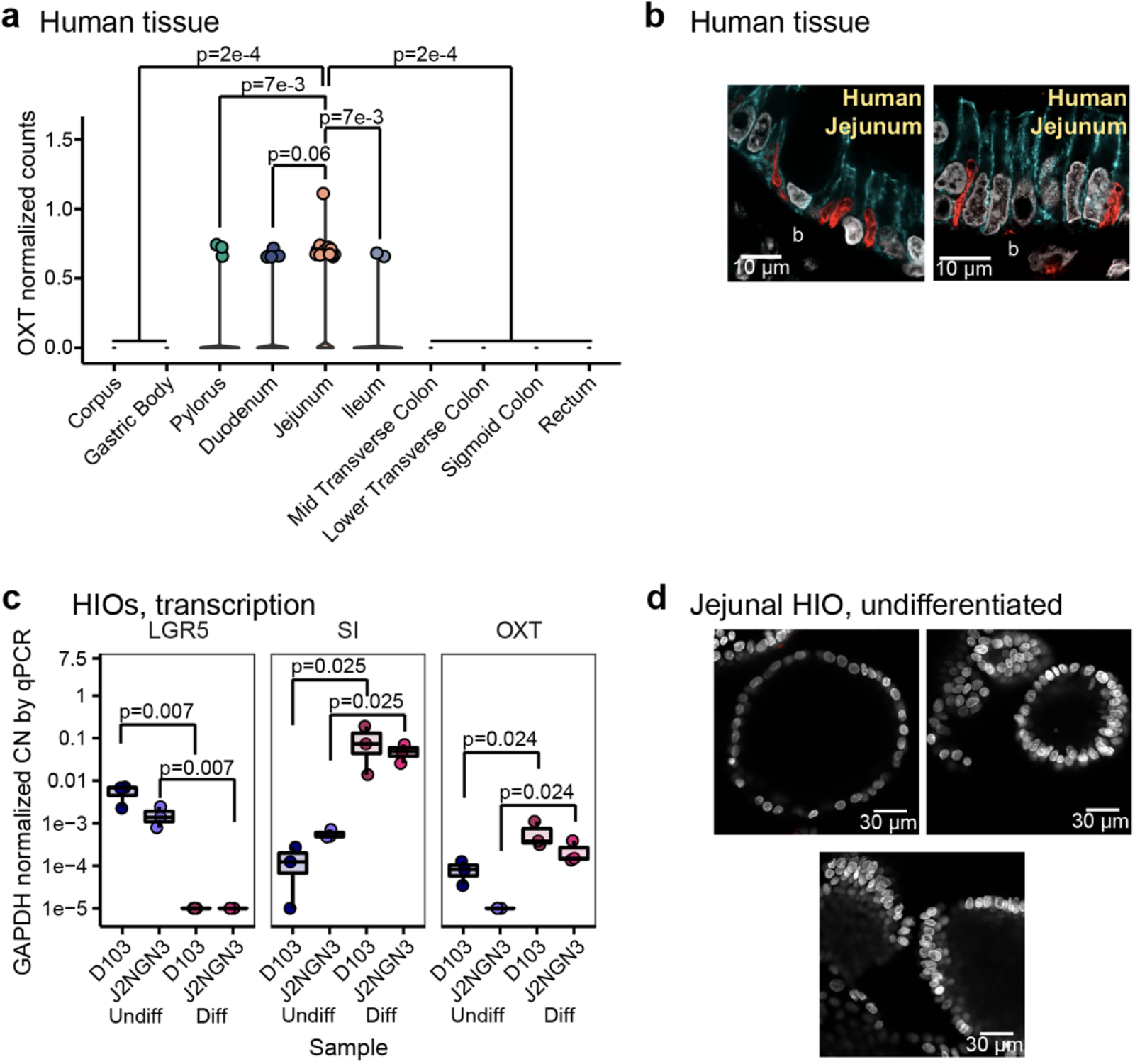
Oxytocin expression and production in the epithelium of the human gastrointestinal tract. **a)** Log normalized counts of oxytocin expression in various human intestinal epithelial tissues reported by the scRNA-Seq data of the Human Cell Landscape^39^. Significance values reflect the number of rarefactions (of 10,000) in which the comparison had a *p* value >0.05 by a Dunn Test with a Benjamini-Hochberg correction. These *p* values were similar whether the number of cells expressing oxytocin or oxytocin expression counts were used. Only *p* values <0.05 are shown. **b)** Oxytocin visualized by immunofluorescence imaging in 35 μm sectioned human jejunum. **c)** Copy number (CN) of *LGR5* (stem cell marker), *SI* (differentiation marker), and *OXT* transcripts by rt-qPCR, normalized to *GAPDH* CN in undifferentiated and differentiated duodenal (D103) and jejunal (J2-*NGN3*, differentiated but not induced to increase enteroendocrine cells) organoids. Dots represent averaged triplicate qPCR data each from a separate pooled batch of three 3D organoid wells. Where product could not be amplified during rt-qPCR, a *GAPDH* CN normalized value of 1e-5 was used. Significance values were determined from the least squares means derived from a linear model with pairwise comparisons corrected using a Benjamini-Hochberg multiple testing correction (see **Supplementary Tables 2** and **3**). **d)** Oxytocin visualized by confocal immunofluorescence in undifferentiated 3D J2-*NGN3* organoids. In the images, DAPI stained nuclei are shown in white, oxytocin staining in red, and E-cadherin staining in cyan. Basolateral (b) sides are labeled. Undiff, undifferentiated. Diff, differentiated. a: corpus: *n* = 1,498 cells, gastric body: *n* = 6,161 cells, pylorus: *n* = 1,281 cells, duodenum: *n* = 1,721 cells, jejunum: *n* = 3,508 cells, ileum: *n* = 1,303 cells, epityphlon: *n* = 967 cells, ascending colon: *n* = 232 cells, mid transverse colon: *n* = 3,192 cells, lower transverse colon: *n* = 4,698 cells, sigmoid colon: *n* = 1,724 cells, rectum: *n* = 3,464 cells, each region was derived from a single patient; c: *n* = 3 experiments. **Extended Data Movie: Oxytocin visualized by immunofluorescence in differentiated 3D J2-*NGN3* organoids**. White are DAPI stained nuclei, oxytocin in red, and E-cadherin staining in cyan. Movie constructed from z-stack confocal images.

**Extended Data Fig. 2:**
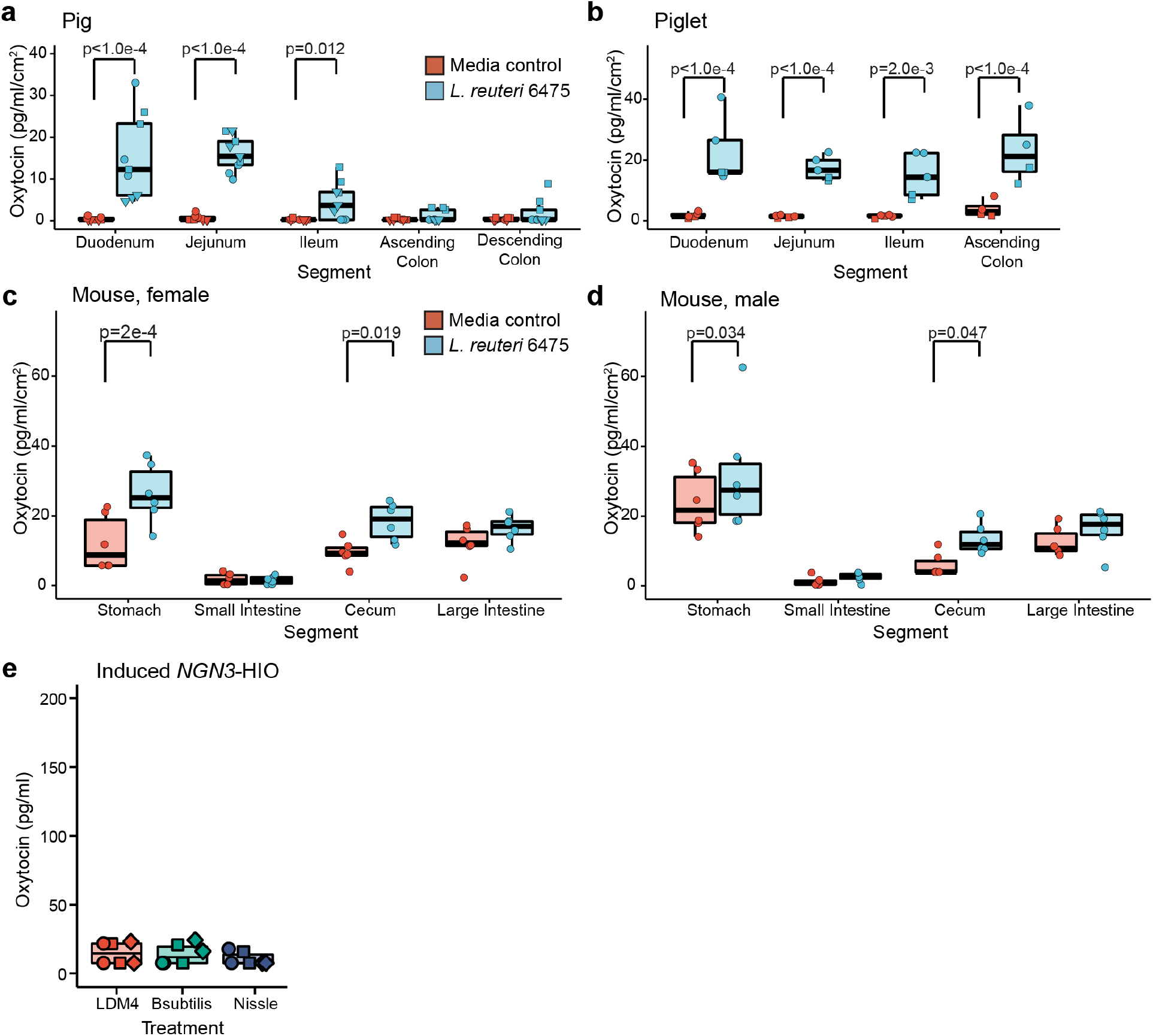
Oxytocin secreted from pig, piglet, and mouse intestinal tissue in response to *L. reuteri*-conditioned medium. Oxytocin measured by ELISA and normalized by tissue surface area secreted from *ex vivo* **a)** pig, **b)** piglet, **c)** female mouse, and **d)** male mouse intestinal tissue. Point shapes indicate unique animal (in a and b). Each point is a unique animal in c and d. Treatment groups are colored as indicated in a and c. Data for the jejunum for a and b are also shown in Fig. 2c and d and data from the cecum for c and d are also shown in Fig. 2e. **e)** Oxytocin measured by ELISA from induced J2-*NGN3* HIOs treated with bacterial growth medium (LDM4) or bacterial-conditioned medium from *Bacillus subtilis* or *Escherichia coli* Nissle. Data presented were generated with those in Fig. 2f, and the LDM4 data shown are the same in both figures. Point shape reflects HIO batch. Significance values were determined as in Fig. 2. a: *n* = 3 animals per region and condition with three replicate tissues (9 datapoints total); b: *n* = 2 animals per region and condition with three replicate tissues (6 datapoints total); c, d: *n* = 6 animals per region and condition; e: *n* = 3 HIO batches with two replicate monolayers per condition (6 datapoints total).

**Extended Data Fig. 3:**
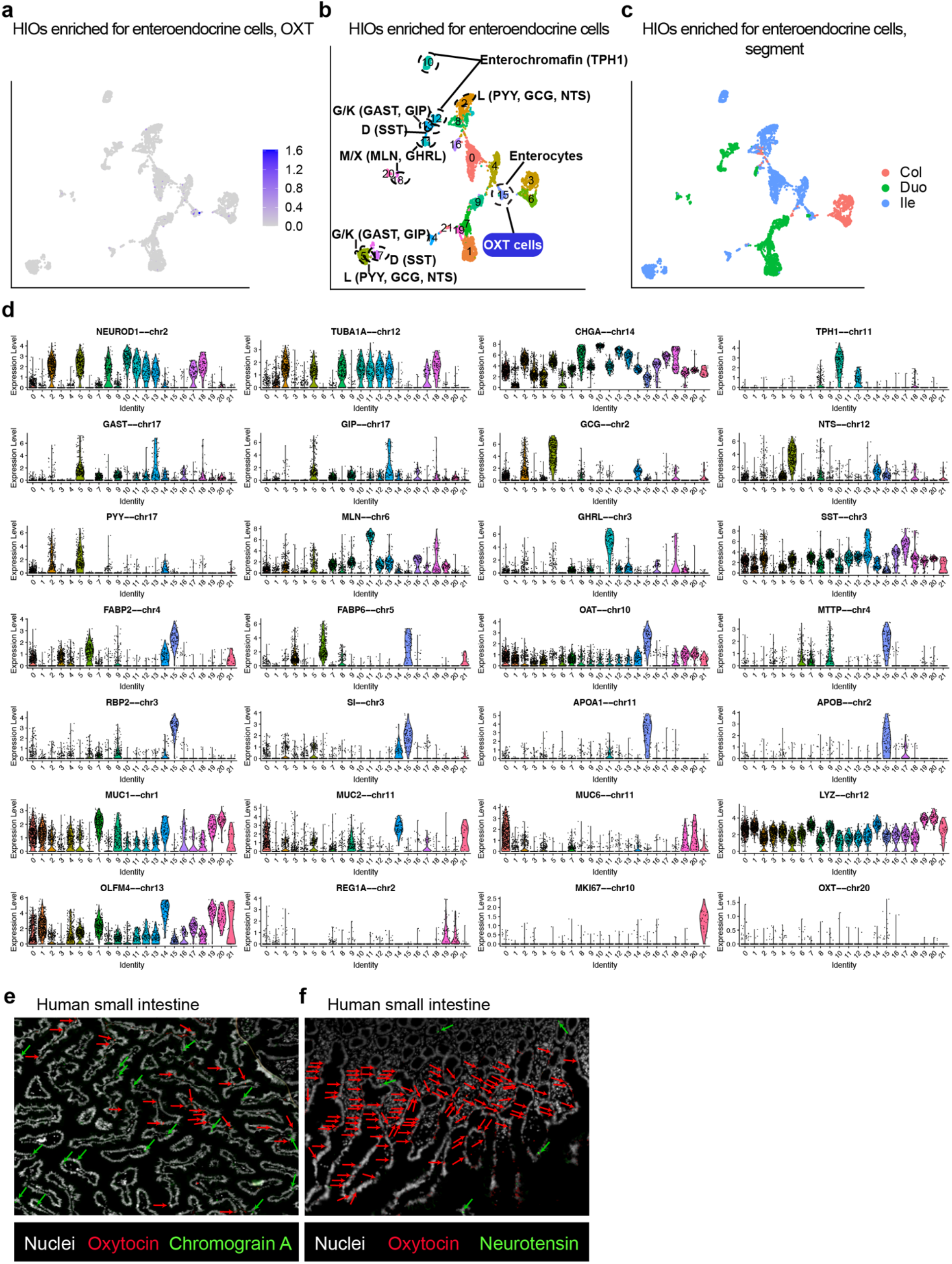
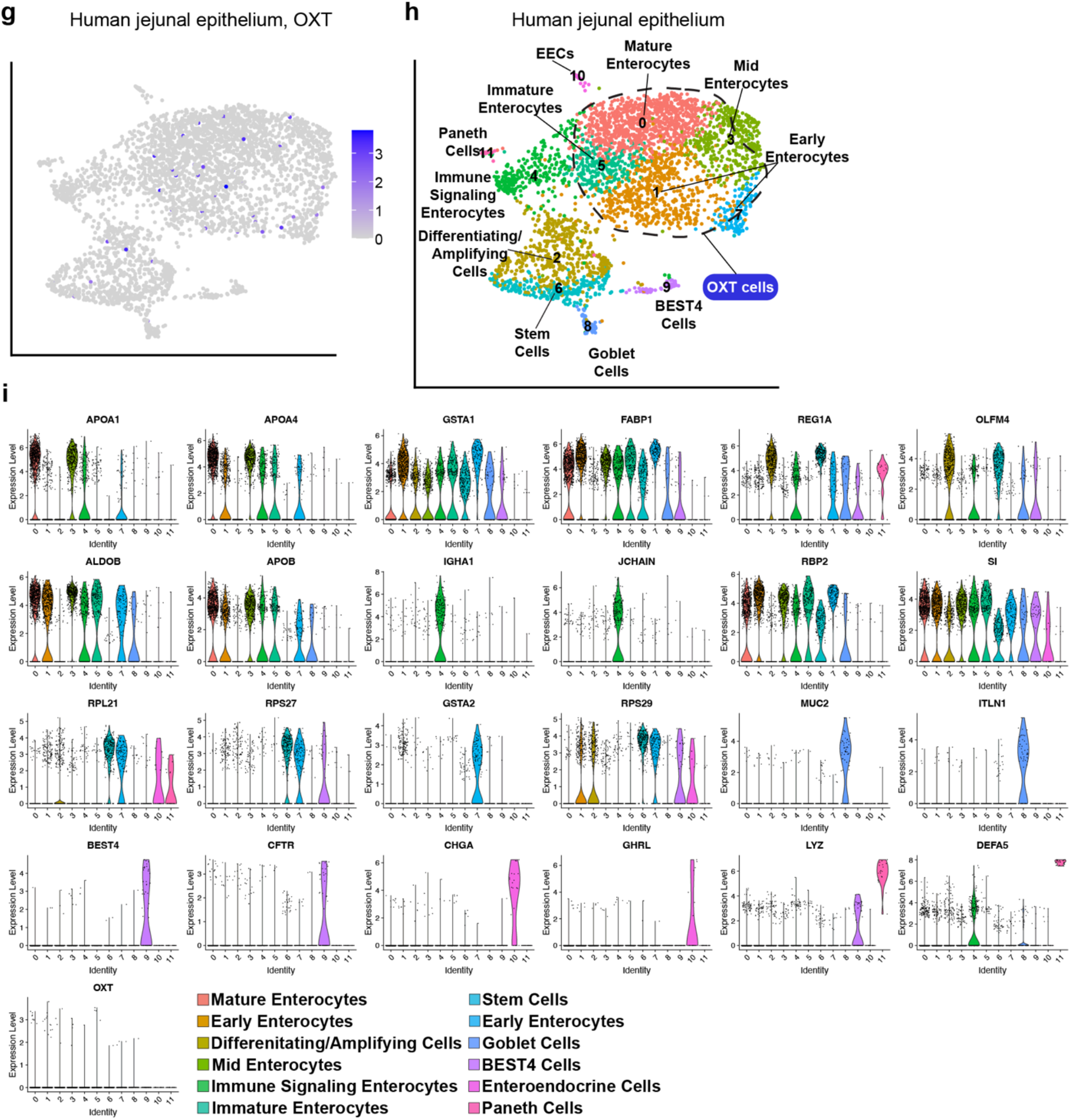
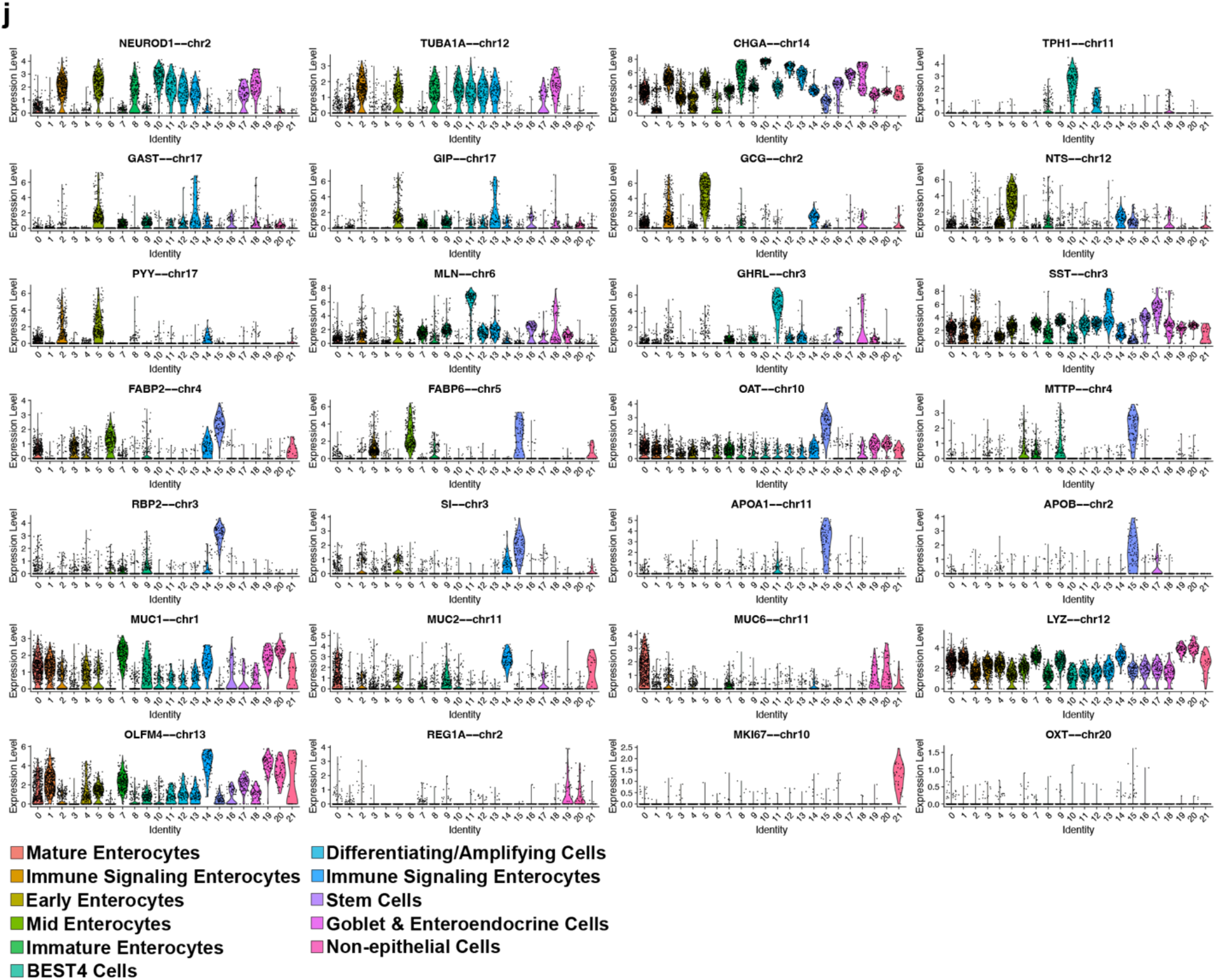
Oxytocin transcription in enterocytes. UMAP of Beumer et al 2020^46^ scRNA-Seq data labeled with **a)** oxytocin counts, **b)** identified cell clusters, or **c)** colored by intestinal segment. **d)** Gene enrichments used to annotate clusters in b. Also see **Supplementary Table 4**. Oxytocin (red) and **e)** chromogranin A (green) or **f)** neurotensin (green) labeled in 6 μm sectioned human small intestinal tissue. Nuclei stained with DAPI are shown in white. UMAP of Human Cell Landscape scRNA-Seq data^39^ labeled with **g)** oxytocin counts or **h)** identified cell clusters. **i)** Gene enrichments used to annotate clusters in h. Also see **Supplementary Table 5. j)** Gene enrichments used to annotate clusters in **Fig. 3d**. Also see **Supplementary Table 6**. a-d: *n* = 4,383 cells from 19 different combinations of cell line and treatment; g-i: *n* = 3,508 cells from a single patient; j: *n* = 2,791 cells from 4 different patients.

**Extended Data Fig. 4:**
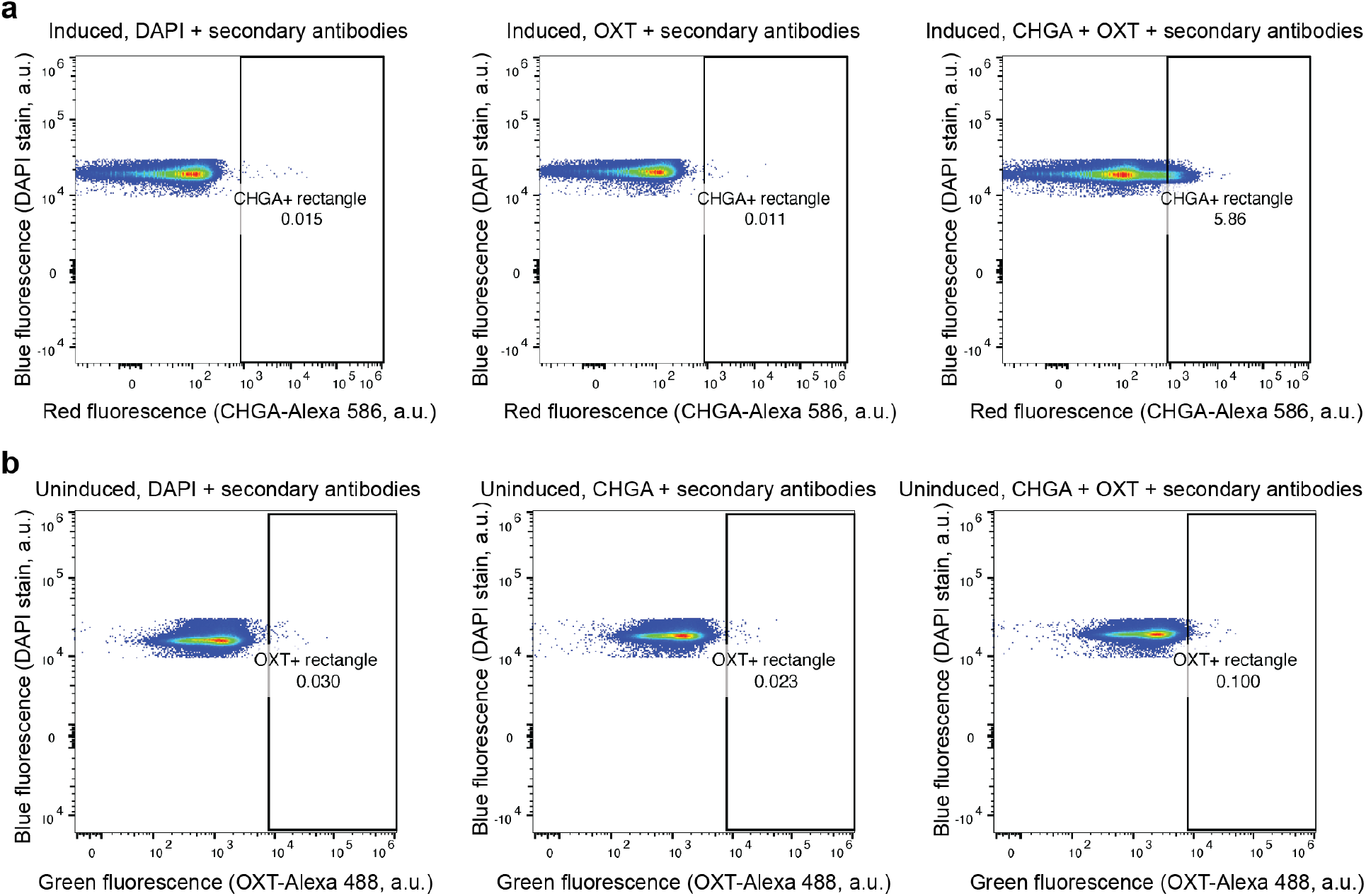
Gating strategy for flow cytometry. Example gating strategies for **a)** CHGA and **b)** OXT. For uninduced and induced J2-*NGN3* HIOs separately, gating for CHGA or OXT was established by looking at all staining controls not containing the CHGA or OXT primary antibody and setting the gate such that ≤ 0.05% positive cells were observed in the relevant channel. In both a and b, the left and middle plots are examples of staining controls and the right most plots are fully stained samples used for quantification. Cells shown are those after selecting for single cells and those in G1.

## Notes

### Competing Interest Statement

Funding was provided by BioGaia (to RAB), with whom we have a provisional patent (HAD, RAB, SCD).

