## Supplementary Tables 1, 8, and 9 for "Microbial stimulation of oxytocin release from the intestinal epithelium via secretin signaling"

**Supplementary Table 1: scRNA-Seq datasets surveyed for *OXT* expression.**

| Dataset | Species | Age group | Type | Intestinal region(s) | Total OXT+ cells | Total cells |
|---|---|---|---|---|---|---|
| Grün et al. Nature 2015 | Mouse | Adult | Organoids | Small intestine | 1 | 1680 |
| Haber et al. Nature 2017 | Mouse | Adult | Epithelial tissue and organoids | Small intestine | 11 | 51897 |
| Beumer et al Nature Cell Biology 2018 | Mouse | Adult | Organoids | Small intestine | 6 | 3072 |
| Fujii et al Cell Stem Cell 2018 | Human | Adult | Epithelial tissue and organoids | Ileal | 29 | 9281 |
| Beumer et all Cell 2020 | Human | Adult | Organoids | Small & large intestine | 125 | 8448 |
| Han et al Nature 2020 | Human | Adult | Whole tissue | Stomach, small, & large intestine | 52* | 29700 |
| Fawkner-Corbett et al Cell 2021 | Human | Fetal | Whole tissue | Small & large intestine | 3082* | 63385 |
| Elmentaite et al Nature 2021 | Human | Fetal, adult | Whole tissue | Small & large intestine | 8523* | 142113 |
| Li et al. Cell Regeneration 2022 | Mouse | Adult | Epithelial tissue | Ileum | 2 | 1144 |
| Li et al. Cell Regeneration 2022 | Rat | Adult | Epithelial tissue | Ileum | 0 | 3047 |
| Li et al. Cell Regeneration 2022 | Pig | Adult | Epithelial tissue | Ileum | 0 | 913 |
| Li et al. Cell Regeneration 2022 | Macaque | Adult | Epithelial tissue | Ileum | 220 | 891 |

*count post-filtering for epithelial cells

**Supplementary Table 2: Statistical models.** Provided in Excel document.

**Supplementary Table 3: Effect sizes.** Provided in Excel document.

**Supplementary Table 4: Genes differentially expressed among clusters permitting cluster annotation in the Beumer et al 2020 dataset.** Provided in Excel document.

**Supplementary Table 5: Genes differentially expressed among clusters permitting cluster annotation in the jejunum epithelium Human Cell Landscape dataset**. Provided in Excel document.

**Supplementary Table 6: Genes differentially expressed among clusters permitting cluster annotation in the jejunum Gut Cell Atlas dataset.** Provided in Excel document.

**Supplementary Table 7: Genes differentially expressed between cells expressing *OXT* and not in the jejunum Gut Cell Atlas dataset.** Provided in Excel document.

### Supplementary Table 8: Antibodies used for imaging and flow cytometry.

| Antibody type | Target | Host | Company | Catalog # | Dilution |
|---|---|---|---|---|---|
| Primary | Chromogranin A | Mouse | Santa Cruz, USA | sc-393941 | 1:100 (6 µm tissue); 1:50 (flow cytometry) |
| Primary | Neurotensin | Mouse | Santa Cruz, USA | sc-377503 | 1:100 (6 µm tissue) |
| Primary | Oxytocin | Rabbit | Sigma-Aldrich, USA | HPA071892 | 1:100 (3D organoids) |
| Primary | Oxytocin | Rabbit | ImmunoStar, USA | 20068 | 1:500 (6 µm tissue); 1:100 (35 µm tissue); 1:4000 (3D organoids); 1:100 (flow cytometry) |
| Primary | Sucrase isomaltase | Mouse | Santa Cruz, USA | sc-393470 | 1:25 (6 µm tissue) |
| Primary | MTP | Mouse | Santa Cruz, USA | sc-515742 | 1:100 (6 µm tissue) |
| Primary | OAT | Mouse | Santa Cruz, USA | sc-374243 | 1:25 (6 µm tissue) |
| Primary | APOA1 | Mouse | Santa Cruz, USA | sc-376818 | 1:25 (6 µm tissue) |
| Primary | Aldolase B | Mouse | Santa Cruz, USA | sc-393278 | 1:25 (6 µm tissue) |
| Secondary | Anti-mouse Alexa Fluor 488 | Goat | Life Technologies, USA | A-11001 | 1:300 (6 µm tissue) |
| Secondary | Anti-mouse Alexa Fluor 568 | Goat | Invitrogen, USA | A-11004 | 1:600 (flow cytometry) |
| Secondary | Anti-rabbit Alexa Fluor 488 | Goat | Invitrogen, USA | ab150077 | 1:800 (3D organoids and flow cytometry) |
| Secondary | Anti-rabbit Rhodamine Red-X | Goat | Jackson ImmunoResearch, USA | AB_2338028 | 1:200 (6 µm tissue) |
| Primary-Conjugated | Alexa Fluor 647 conjugated E-cadherin | Mouse | BD Pharmingen, USA | 560062 | 1:50 (6 µm tissue); 1:10 (35 µm tissue) |
| Stain | NucBlue Fixed Cell Stain | NA | Invitrogen, USA | R37606 | 0.07 to 0.1x (6 µm tissue); undiluted (35 µm tissue); 0.07x (3D organoids); 0.07x (flow cytometry) |

### Supplementary Table 9: Primers for rt-qPCR.

| Target | Forward Sequence | Reverse Sequence | Purpose | Citation |
|---|---|---|---|---|
| *OXT* | GCTGAAACTTGATGGCTCCG | TTCTGGGGTGGCTATGGG | Detect *OXT* | Wang et al. 2008. *Molecular Psychiatry.* **13**: 786-799. |
| *LGR5* | CTCCCAGGTCTGGTGTGTTG | GAGGTCTAGGTAGGAGGTGAAG | Marker for stem cells | Chang-Graham et al. 2019. *CMGH*. **8**: 209-229. |
| *SI* | CATCCTACCATGTCAAGAGCCAG | GCTTGTTAAGGTGGTCTGGTTTAAATT | Marker for enterocytes | Sclafani et al. 2007. *Proc Natl Acad Sci USA*. 104:14887-14888. |
| *GAPDH* | ACCACAGTCCATGCCATCAC | TCCACCACCCTGTTGCTGTA | qPCR normalization | Wang et al. 2015. *Cell & Bioscience*. **5**:3. |
| *CHGA* | TGTAGTGCTGAACCCCCACC | CTCTCGCCTTTCCGGATCT | Marker for enteroendocrine cells | Chang-Graham et al. 2019. *CMGH*. **8**:209-229. |
